# Functional Connectivity Alterations in Cocaine Use Disorder: Insights from the Triple Network Model and the Addictions Neuroclinical Assessment Framework

**DOI:** 10.1101/2024.11.12.623073

**Authors:** Ziyang Xu, Lie Li, Ruobing Liu, Mohamed Azzam, Shibiao Wan, Jieqiong Wang

## Abstract

Cocaine use disorder (CUD) disrupts functional connectivity within key brain networks, specifically the default mode network (DMN), salience network (SN), and central executive network (CEN). While the triple network model has been proposed to explain various psychiatric disorders, its applicability to CUD requires further exploration. In the present study, we built machine learning classifiers based on different combinations of DMN/SN/CEN to distinguish cocaine-use disorder (CUD) subjects from healthy control (HC) subjects. Among them, the combination of the SN and the CEN results in a remarkably high accuracy of 73.4% (sensitivity/specificity: 69.6%/78.6%, AUC: 0.78), outperforming the model based on the full triple network. This supports the hypothesis that during the binge/intoxication stage of addiction, the SN and the CEN play a more critical role than the DMN, consistent with the Addictions Neuroclinical Assessment (ANA) framework. Functional connectivity analysis revealed decreased connectivity within the DMN and the SN and increased connectivity within the CEN in CUD patients, suggesting that alterations in these networks could serve as biomarkers for addiction severity.

## 1. Introduction

Cocaine could be used for medical purposes, such as local anesthesia and decreasing bleeding in surgeries. But it is always illegally used as a recreational drug to make people addicted. Patients with cocaine addiction have a high risk of suffering heart attacks, liver/kidney/lung problems, severe depression, etc. The cocaine addiction is related to the increase of the dopamine transferred in the reward system of the brain (Di Chiara & Bassareo, 2007). It is important to understand the neuromechanism of cocaine addiction to help people get off cocaine.

The brain is known to be affected by cocaine for years. The gray matter concentration/volume is decreased in the insula (Franklin et al., 2002) and the thalamus (Sim et al., 2007) in patients with cocaine addiction. Cocaine also impairs the white matter integrity of the anterior corpus callosum (Moeller et al., 2005). Recently, more studies have focused on resting-state functional connectivity (FC) in cocaine-use disorder (CUD) since the FC study can explore the interactions between the brain regions or brain networks, resulting in a system-level understanding of the brain (Bressler & Menon, 2010; Van Den Heuvel & Hulshoff Pol, 2010). The results of the early FC study that reduced FC in the primary visual cortex and motor cortex after cocaine administration reflected the changes in neuronal activity (Li et al., 2000). The following whole-brain FC analysis revealed a general decrease in FC between most regions within the mesocorticolimbic (MCL) circuit (including the amygdala, the hippocampus, the ventral anterior cingulate cortex, etc.) and interconnected brain areas, which further provided support for the importance of salience network (SN) in cocaine dependence (Gu et al., 2010). The reduced FC between the inferior frontal sulcus and the lateral prefrontal cortex and parietal areas in cocaine addiction shows that the central executive network (CEN) is involved in cocaine dependence (Kelly et al., 2011). The decreases in the FC between the default-mode network (DMN) and the hippocampus exhibit the importance of the DMN in cocaine (Ding & Lee, 2013).

The DMN, the SN, and the CEN have been proven to be related to many psychiatry and neurological disorders (Banich et al., 2009; Öngür et al., 2010; Stein et al., 2007), which was summarized as a unifying triple network in psychopathology (Menon, 2011). In 2012, Matthew et al. imported the concept of the triple network to the addiction domain, with nicotine abstinence as an example (Sutherland et al., 2012). The later study of the same group developed an index integrating the SN-CEN and SN-DMN correlations and proposed the possibility of this index as a biomarker for smoking cessation (Lerman et al., 2014). Liang et al. revealed the disruption of network-level interactions involving the DMN/SN/CEN in cocaine addiction (Liang et al., 2015). McHugh et al. proved the interhemispheric connectivity between the ECN and SN related to cocaine addiction (McHugh et al., 2017). Geng et al. identified the alternation of functional circuits in the SN and the DMN of addiction to cocaine, emphasizing the importance of these networks in the treatment of cocaine dependence (Geng et al., 2017).

Instead of the triple network framework, the Addictions Neuroclinical Assessment (ANA), a neuroscience-based framework, was proposed for addictive disorders (Kwako et al., 2016). The ANA framework focuses more on three domains—executive function, incentive salience, and negative emotionality—which are crucial in understanding the different stages of addiction (Koob & Volkow, 2016). The ANA framework also identifies three stages of addiction: binge/intoxication, withdrawal/negative effect, and preoccupation/anticipation/craving. Together, these domains and stages form the core functional elements of addictive disorders, providing a comprehensive approach to their research and treatment.

In this study, we will investigate which framework (the triple network or the ANA framework) is more consistent with the neuromechanism of cocaine addiction to help people get off cocaine. We hypothesize that the ANA framework is a better explanation than the triple network during the binge/intoxication stage of cocaine dependence, which means the SN and the CEN play more important roles than the DMN in the binge/intoxication stage of cocaine dependence. To test it, we built machine learning (ML) frameworks based on the triple network to classify the patients with cocaine use disorder (CUD) from the normal controls based on the triple networks

## 2. Materials and methods

### 2.1. Participants

The public-accessible dataset, the SUDMEX_CONN dataset (Angeles-Valdez et al., 2022), was used in this study. This dataset included 74 cocaine-use disorder (CUD) patients (age: 31.0±7.20, Male (M)/Female (F): 65/9) and 64 health control (HC) subjects (age: 30.6 ± 8.26, M/F: 53/11). The magnetic resonance imaging (MRI) data of this dataset includes T1-weighted imaging data, diffusion-weighted (DWI) data, and resting-state functional MRI (rs-fMRI) data. The details can be found in the paper describing this dataset (Angeles-Valdez et al., 2022).

Among them, 35 HC subjects (age: 30.14 ± 7.73, M/F: 27/8) and 48 CUD subjects (age: ± 7.38, M/F: 45/3) were chosen in the study by the following criteria: (1) with both rs-fMRI data and T1-weighted imaging data, (2) do not have other substance use dependence, (3) passing image quality control, (4) with small head motion during the rs-fMRI scan, (5) making the two groups age-sex-matched. The CUD patients and HC subjects are matched in terms of age (two-sample T-test, p=0.190) and sex (chi-square test, p=0.061).

### 2.2. Data preprocessing

The DPARSF (Data Processing Assistant for Resting-State fMRI, http://www.rfmri.org/DPARSF) toolbox (Yan, 2010) was applied to preprocess the resting-state fMRI data. Considering the magnetization equilibration, the first 10 time points of each time series were removed. The remaining fMRI volumes were corrected by slices, and head realignment was conducted. In order to reduce the effect of head motion, the fMRI data with big head motion were excluded, i.e., translational or rotational motion parameters were over 3 mm or 3°. The nuisance covariate effects of the white matter and cerebrospinal fluid, as well as the Friston 24 head-motion parameters, were removed. Subsequently, temporal band-pass filtering (0.01<f<0.1 Hz) was performed, and the fMRI data were normalized to the Montreal Neurological Institute (MNI) space via the segmented results of T1-weighted images. The normalized fMRI data were then resampled to 3 mm × 3 mm × 3 mm voxels. The Gaussian kernel with a full width at half-maximum (FWHM) of 4 mm was used to smooth the data.

### 2.3. Identification of the DMN/SN/CEN

The group information-guided independent component analysis (GIG-ICA) was applied to compute the specific independent component (IC) of each subject with correspondence across all subjects (Du and Fan 2013). Firstly, the tool Melodic (default setting) in FSL was used to compute the group template with 25 ICs based on the preprocessed resting-state fMRI data. Then, the group template was used as a reference to calculate subject-specific ICs (also referred to as intrinsic brain networks, IBNs) and corresponding time series with a multi-objective optimization solver. Then, Yeo’s Atlas (Thomas Yeo et al., 2011) was applied to identify the belong of the DMN, the SN, and the CEN for each IBN.

### 2.4. Machine learning models to classify CUD patients from HC subjects based on the DMN/SN/CEN

The IBNs of these three resting-state networks were used as bases for a linear subspace and were analyzed on the Grassmann manifold (Harris, 1992) to calculate Riemannian distance. Then, the Riemannian distance was used in conjunction with a support vector machine (SVM) to build the classifier (Fan et al., 2010, 2011). A 10-fold cross-validation (CV) was used to evaluate the performance of each of the classifiers. Specifically, all subjects were randomly divided into 10 subsets with almost equal size. These 10 subsets were used in 10 training-testing runs. In each run, one of the 10 subsets was used as the testing set, and the other nine subsets were used as the training set. The training-testing runs were repeated until all 10 subsets had been used as the testing set. Then, the accuracy of the classification, the sensitivity, and the specificity were calculated based on the 10 training-testing runs. A receiver operating characteristic (ROC) curve and the area under the curve (AUC) were also computed based on the classification scores of all subjects. In order to avoid statistical bias, the 10-fold CV was repeated 10 times.

Seven classifiers were trained to differentiate cocaine use disorder (CUD) patients from healthy control (HC) subjects using IBN features. These classifiers were based on:

1. IBN features from a single network: the DMN, the SN, or the CEN.
2. IBN features from combinations of two networks: DMN-SN, DMN-CEN, and SN-CEN.
3. IBN features from all three networks combined: DMN, SN, and CEN.

### 2.5. Statistical analysis

The voxel-wise functional connectivity (FC) map of each IBN in the DMN/SN/CEN was obtained by computing the Pearson correlation coefficient between the time course of the IBN and that of each voxel in the gray matter and the following z-score transformation. The two-sample t-test was conducted to compare the functional connectivity (FC) of each IBN between the HC subjects and the CUD patients (voxel p<0.01, cluster p<0.05, two-tailed Gaussian Random Field (GRF) corrected), with age, sex, education level, head motion values and tobacco usage values as covariates. The group comparison of the FC between each pair of IBNs in the Triple Network was also conducted by a two-sample t-test (p < 0.05). In addition, the relationship between classification scores and the clinical measures was measured by Pearson’s correlation analysis within the subjects with CUD. The altered FC within the IBNS or between the IBNs was also correlated with clinical measures in the CUD subjects group. The significant threshold was set at 0.05.

## 3. Results

### 3.1. DMN/SN/CEN Identification

With the GIG-ICA method, the brain was divided into 25 IBNs (Figure 1). The DMN, the SN, and the CEN were identified based on Yeo’s atlas (Thomas Yeo et al., 2011). Among 25 IBNs, IBN 1, IBN 8, and IBN 12 belonged to the DMN. The salience network included IBN 14, IBN 18, and IBN 21. IBN 17 and IBN 23 both belonged to the CEN.

**Figure 1.**
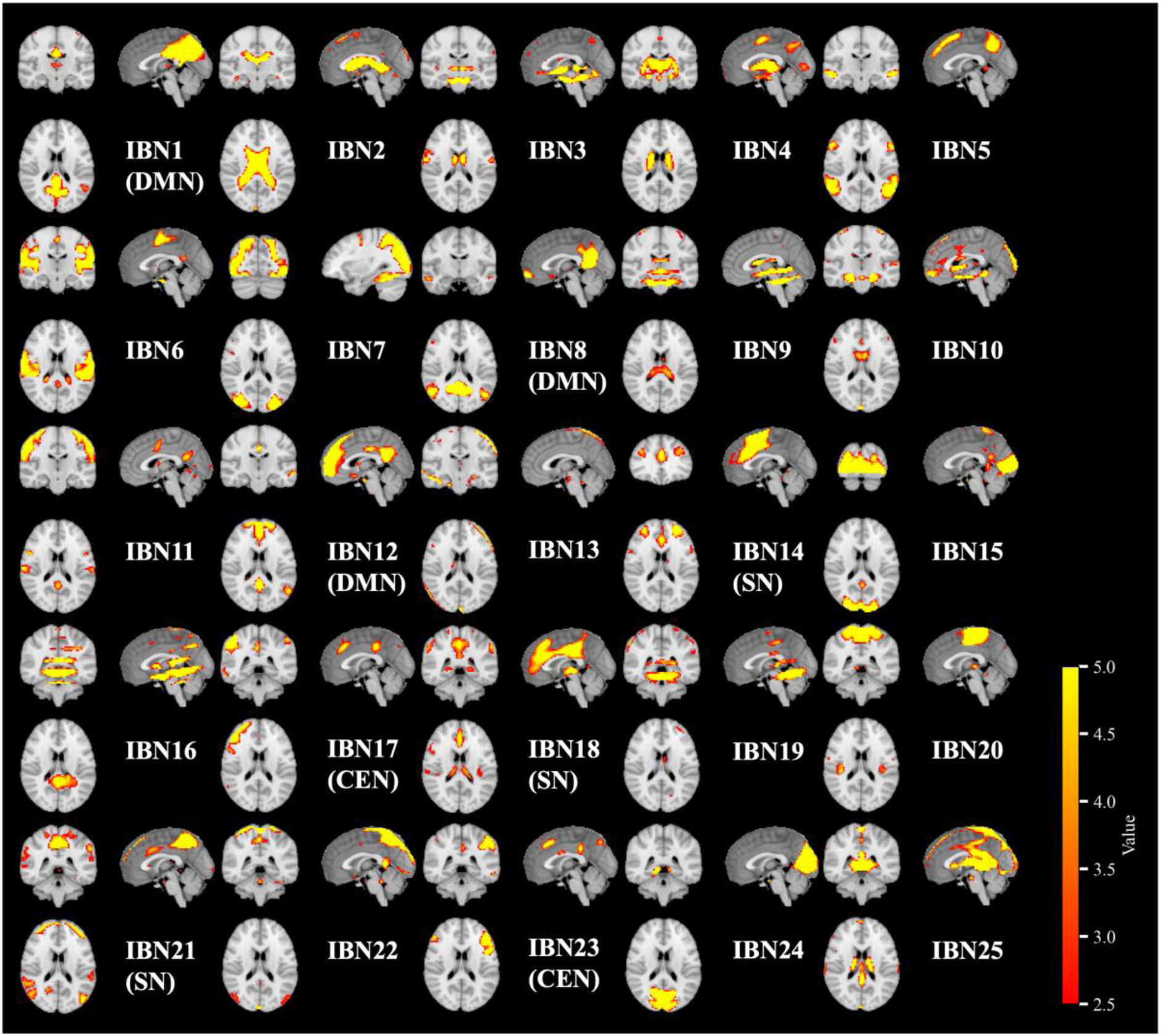
Twenty-five spatial maps of intrinsic brain networks obtained by GIG-ICA. Spatial maps were shown by a one-sample t-test of independent components (z-score maps) of all subjects. IBN: intrinsic brain networks, CEN: central executive network, DMN: default mode network, SN: salience network.

### 3.2. Performance of ML models

The SVM classification models were built on IBNs from the single network, the two-network combination, and the triple-network combination to classify CUD subjects and HC subjects. For the classifiers trained with IBNs from single networks, the one trained with the SN has the best performance compared with the ones trained with the other two networks, with an accuracy of 69.3%±3.4%, sensitivity of 68.3%±4.6%, specificity of 70.6%±3.3%, and AUC of 0.733±0.018. The classifier trained solely on the DMN performed the worst among all classifiers. The classifier trained with IBNs from the combination of SN and CEN got the most distinguished performance among all the combinations of the Triple Network, even outperforming the combination of all three networks, obtaining an accuracy of 73.4%±2.6%, the sensitivity of 69.6%±2.2%, and the specificity of 78.6%±4.3%. The area under the receiver operating characteristic (ROC) curve (AUC) was 0.78±0.02 (Table 1 and Figure 2). The performance of the classifier trained on the SN-CEN is even better than that of the classifier trained on the triple networks. These results are consistent with the assumption that during the binge/intoxication stage of addiction, DMN may not be as important as the other two networks, which is consistent with the ANA circuit.

**Figure 2.**
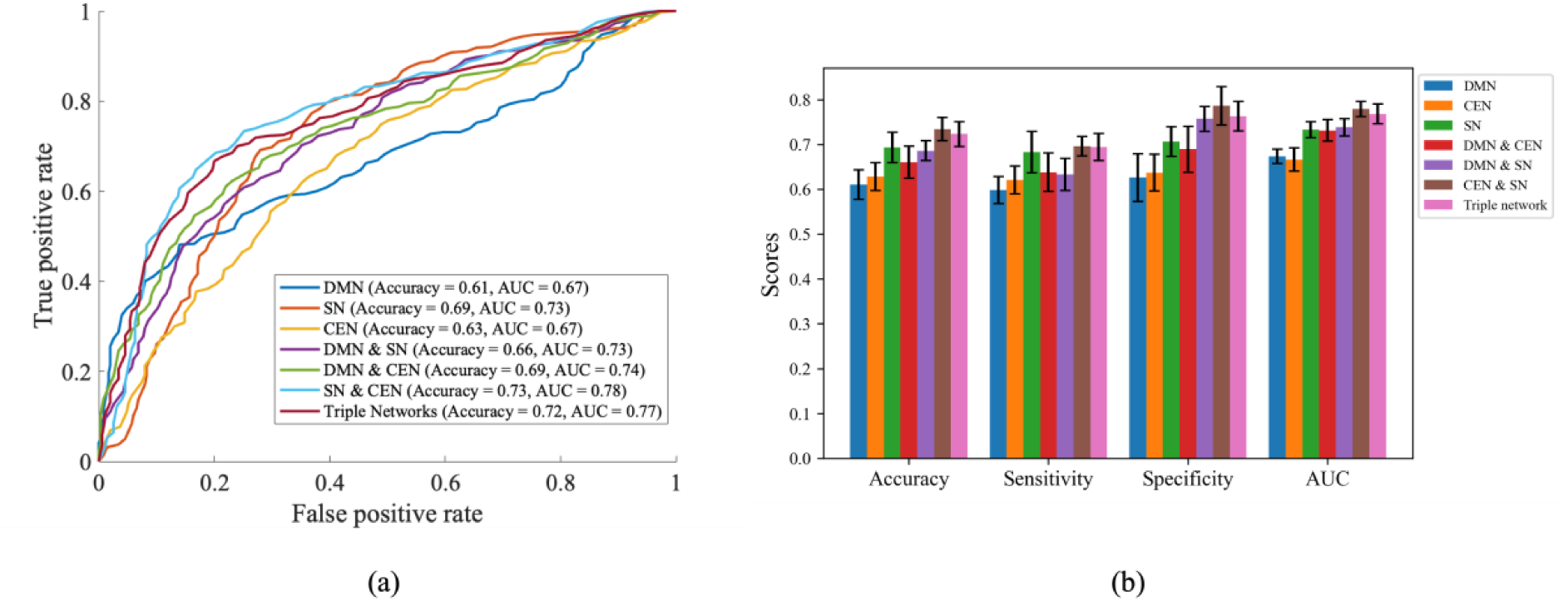
(a) Receiver Operating Characteristic (ROC) Curves of machine learning models based on different combinations of the Triple Network. The ROC curves are averaged over the 10-times 10-fold cross-validation separately (b) Bar plot showing machine learning performance of different classifiers trained on each combination of DMN, SN, and CEN

**Table 1.**
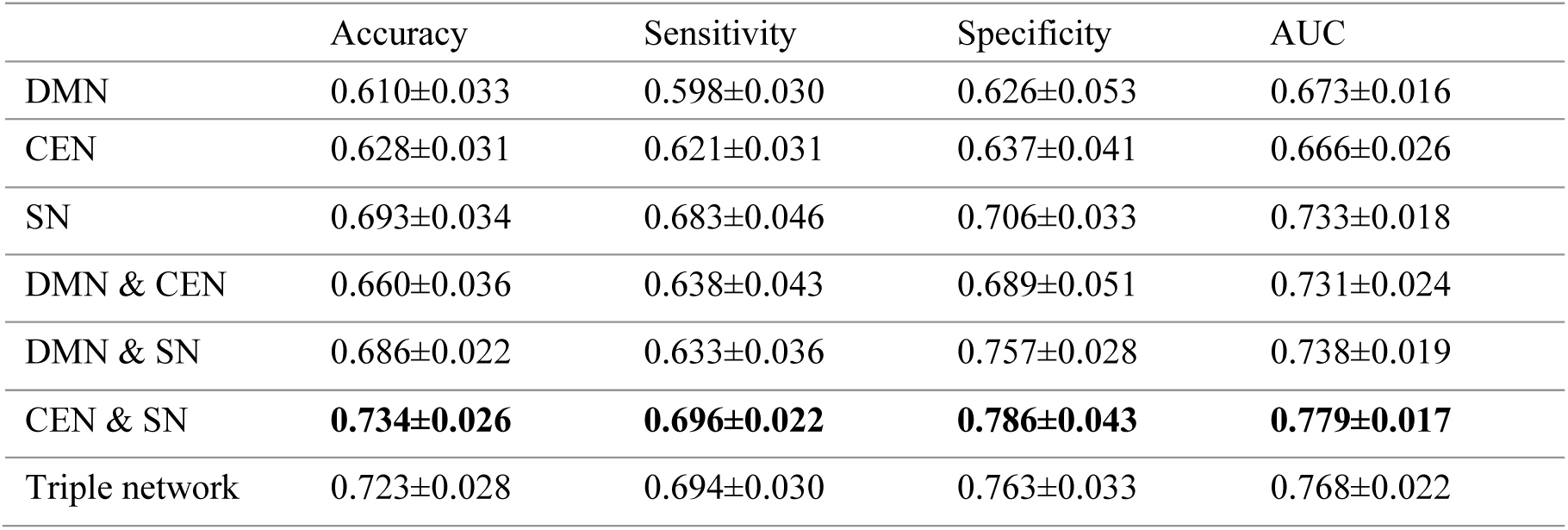
The performance of classifiers trained with different combinations of the triple network.

### 3.3. Abnormal FC between CUD and HC subjects

The whole brain voxel-wise FC measures of the DMN/SN/CEN were compared between CUD patients and HC subjects, and the results are shown in Figure 3 and Table S1. After Gaussian Random Field correction, increased FC (red region in Figure 3) was found within the CEN in the right superior frontal gyrus. while decreased FC (blue regions in Figure 3) was found within the DMN and the SN in the CUD subjects, including the right middle temporal gyrus, the left calcarine fissure, the left middle temporal gyrus, the right superior medial frontal gyrus, the left superior temporal pole, the right precuneus, the right supramarginal gyrus, the right superior temporal pole, the left inferior parietal lobule, and the left insular cortex. The FC measures among the DMN/SN/CEN were also compared between the two groups. (Figure 3).

**Figure 3.**
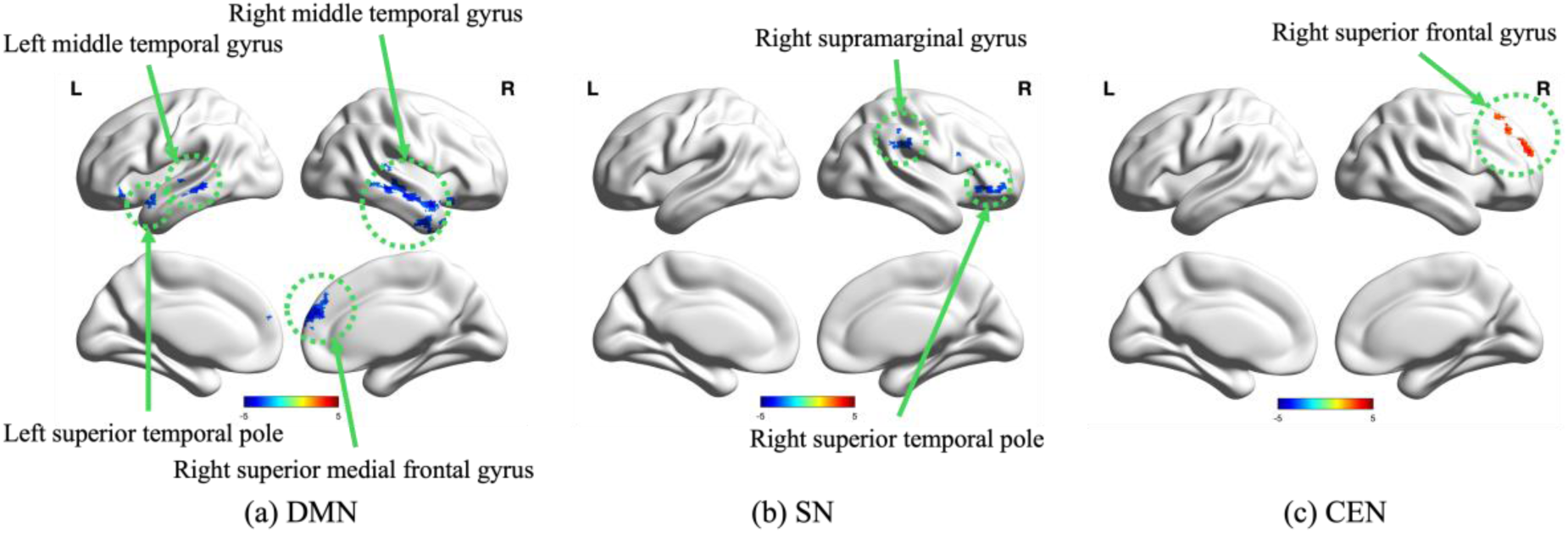
Alterations of functional connectivity (FC) within IBNs of the Triple Network in the CUD patients compared with HC subjects (voxel p<0.01, cluster p<0.05, two-tailed Gaussian Random Field (GRF) corrected). (a) FC alterations within IBNs of the DMN, blue areas represent the decrease in CUD patients; (b) FC alterations within IBNs of the SN, blue areas represent the decrease in CUD patients; (c) FC alterations within IBNs of the CEN, red areas represent the increase in CUD patients

**Figure 4.**
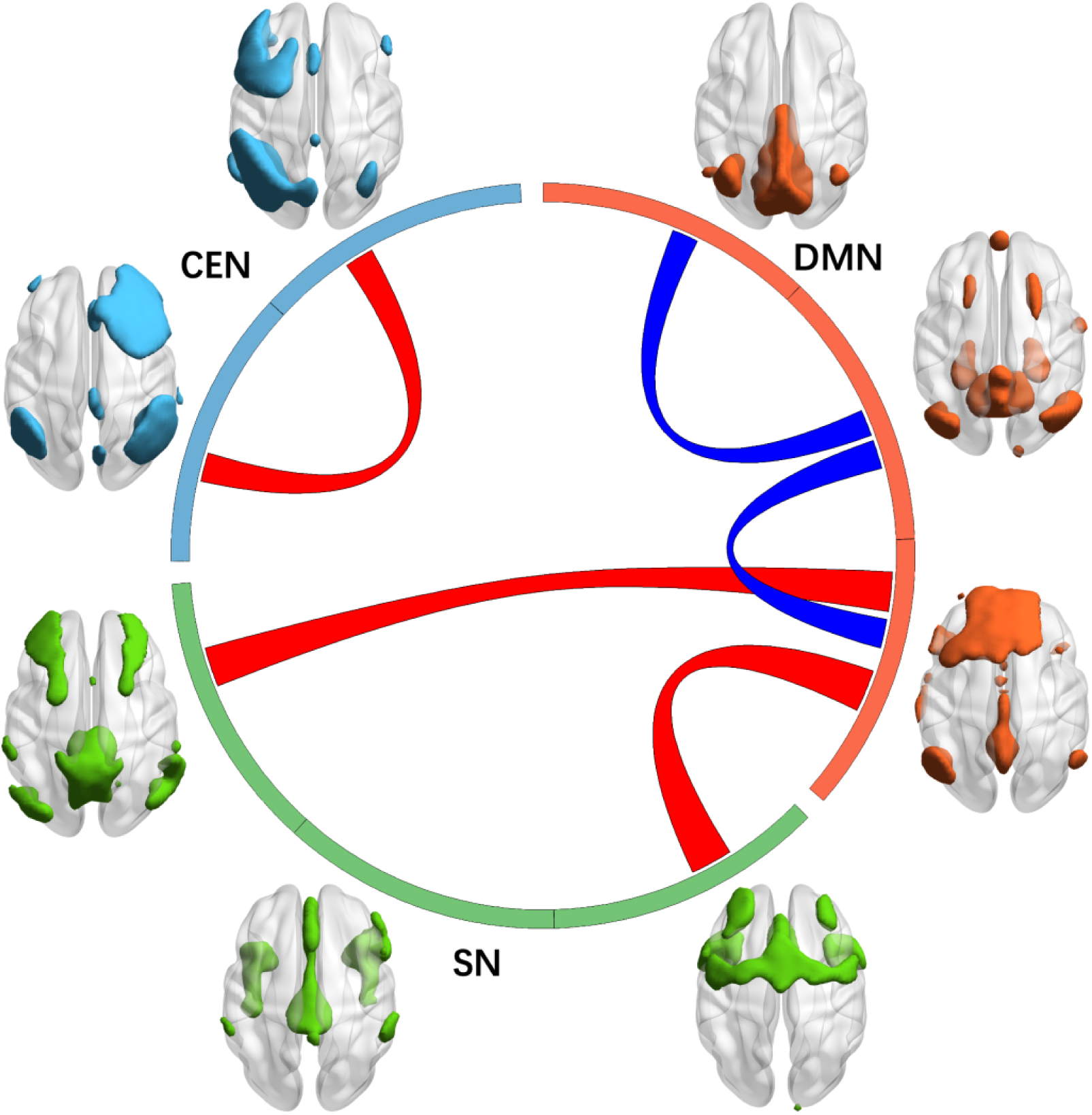
Alterations of functional connectivity (FC) between IBNs of the DMN/SN/CEN in the CUD patients. The red links represent the increased FC (p<0.05), while the blue links represent decreased FC (p<0.05) in CUD subjects. Orange arcs represent DMN, green arcs represent the SN, and light blue arcs represent the CEN.

The FC of each IBN of the triple networks is also compared between CUD patients and HC people. Decreases in FC were found between IBN 1 and IBN 8 as well as between IBN 8 and IBN 12; all these three IBNs belong to the DMN. An increase in FC was found between IBN 17 and IBN 23, both of which belong to the CEN. Two other increases in FC were found between IBN 12 and IBN 14 and between IBN 12 and IBN 21. These increases both happened between the DMN and the SN.

### 3.4. Relationship with clinical measures in subjects with CUD

In order to explore the clinical significance of the classifier, we correlated the classification score of the classifiers with several clinical measures of CUD in the patient group, including years of CUD, days to the last usage, CUD onset age, and dose per week. Significantly positive correlations were found between the years of CUD and classifier scores trained by the DMN-CEN, the SN-DMN, and the triple networks (Table S2). Significant negative correlations were found between the CUD onset age and classifier scores trained by the DMN, the SN, and the SN-DMN. (Table S2). Also, significant negative correlations were found between the CUD onset age and classifier scores trained by the DMN, the CEN, the DMN-CEN, the DMN-SN, and the triple networks (Table S2). There is no significant correlation between the dose per week and the classifier scores.

Then, the correlation coefficients between the average z-transformed value of each cluster and clinical measures are also calculated in CUD patients. And significant correlation was found with the left inferior parietal lobule in the SN, positive to years of CUD and negative to CUD onset age.

We correlated the altered FC within and between IBNs in CUD patients and their clinical measures. The analysis revealed that there was only one significant negative correlation between the altered FC between two CEN components and the number of days to the last usage of cocaine of the CUD subjects (r = -0.427, p = 0.048). No significant correlations were found for FC within the IBNs of the triple networks.

**Figure 5.**
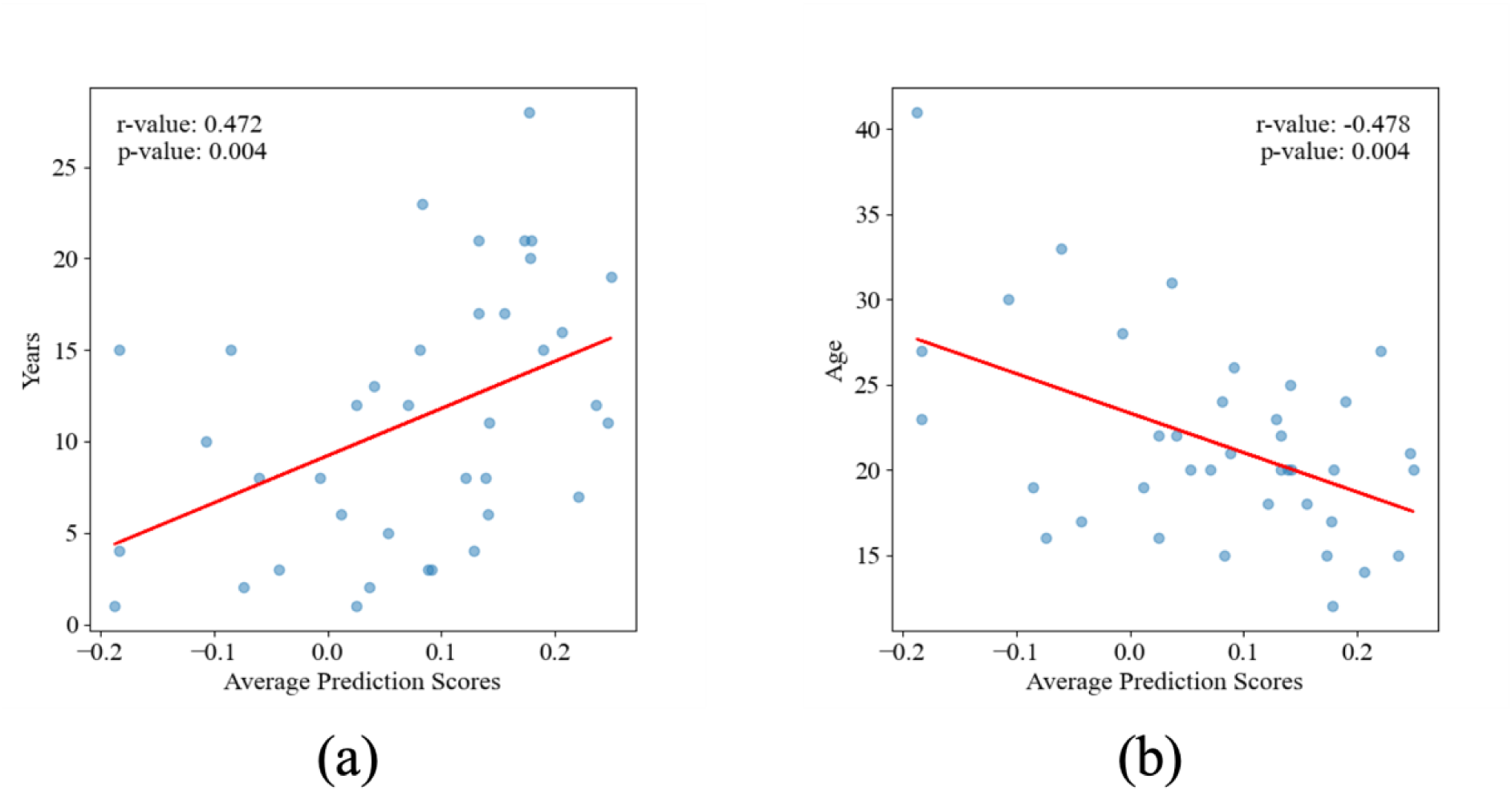
The correlation between the prediction score of the classifier based on the DMN/SN/CEN and the addiction severity index in subjects with cocaine. Each blue dot represents the coordinate corresponding to the average score of 10 predictions for a subject in the classifier and the clinical measure. The red line is the regression curve. (a) A significant positive correlation was found between the years of CUD and the average prediction scores. (b) A significant negative correlation was found between the onset age of CUD and the average prediction scores.

## 4. Discussion

This study applied a data-driven method to build seven classifiers with high classification performance based on the DMN, the SN, and the CEN, as well as the combinations of each two of them and all of the triple networks. Then, functional connectivity is calculated within the triple networks to compare between health control subjects and cocaine-use disorder patients.

Among all the classifiers trained with different combinations of the triple networks, the classifier based on the SN and CEN achieved the highest accuracy of 73.4%±2.6% and an outstanding AUC of 0.779±0.017. The performance of the SN-CEN-based classifier is even better than that of the classifier trained with all the triple networks. The promising performance of distinguishing subjects with CUD from HC subjects at an individual subject level showed that the SN and the CEN play more important roles in the CUD than the triple network model, which is consistent with the Addictions Neuroclinical Assessment (ANA) circuit at the binge/intoxication stage. Also, the classifier trained only with the DMN has the worst performance among all the classifiers, which is also consistent with the study that the DMN has a withdrawal during the binge/intoxication stage.

When compared with HC subjects, the functional connectivity within the triple networks is reduced in the CUD patients in the DMN and the SN but increased in the CUD patients in the CEN. The FC strength was weakened within the DMN for CUD patients, while it was strengthened between the DMN and the SN and within the CEN. These results also supported the hypothesis of the Addictions Neuroclinical Assessment (ANA) circuit that at the binge/intoxication stage, the SN triggers the CEN to be more activated while the DMN is deactivated.

Addiction is strongly correlated with the age of abuse onset, with earlier onset during adolescence increasing the likelihood of stronger addictive tendencies in adulthood (Chen et al., 2009; Grant et al., 2006; Palmer et al., 2009). Additionally, the DMN is considered a valuable biomarker for predicting addiction risk (Zhang & Volkow, 2019), while the CEN is believed to play a crucial role in the treatment of cocaine addiction (McHugh et al., 2017). The scores of all of our machine learning classifiers trained with IBNs from the DMN or the CEN have a significant negative correlation with the CUD onset age and a significant positive correlation with years of CUD. Acute drug or alcohol use has been shown to rapidly disrupt the DMN and the SN (Gorka et al., 2018; Zhang & Volkow, 2019). This aligns with our findings, where the scores from machine learning classifiers trained using IBNs from the DMN, the SN, and their combined DMN-SN network show a significant negative correlation with the number of days since the last usage. All these findings suggest that our models have promising potential for both predicting and aiding in the treatment of cocaine addiction.

The SN was thought to have a core role in the triple network model (Menon, 2011). Signals in the SN triggered the other networks to generate behavioral responses to salient stimuli (Hamilton et al., 2011; Uddin et al., 2011). Our results showed that the classifier trained only with the SN performed better than the other two trained only with the DMN or the CEN, supporting the core role of the SN within the triple networks. Reduced functional connectivity in the SN is believed to lead to the desire for addictive substances, such as nicotine (Sutherland et al., 2012). This was also consistent with our findings of FC decrease within the SN.

Neural changes in the CEN are connected to reward-related decision-making (Yoo et al., 2020). When the reward pathway is activated by immediate rewards, the CEN requires more cognitive resources for less impulsive decisions (McClure et al., 2004). Therefore, chronic substance use disorder can lead to functional connectivity abnormalities in the CEN (Krmpotich et al., 2013; Tapert et al., 2007). Our findings of increased FC within and between IBNs of the CEN were consistent with these reports.

It has been reported that the blood flow was reduced under the psilocybin administration (Carhart-Harris et al., 2012), and drug addictions such as lysergic acid diethylamide (Carhart-Harris et al., 2016) and heroin (Ma et al., 2011) desynchronize brain activity within the DMN. It is also reported that during the intoxication stage, the prominence of the DMN appears to be temporarily decreased (Zhang & Volkow, 2019). Our results showing that the classifier trained on the DMN had the worst performance and that the FC was reduced within and in between the DMN IBNs in CUD subjects were consistent with these reports.

Some limitations should be taken into consideration when interpreting the results. First of all, even though the sexes of the subjects in our experiments are matched, only 8 HC and 3 CUD subjects are female, and most of the subjects are male. The sex has an effect on the behavioral response and the treatment response (Becker et al., 2001; Kosten et al., 1993; Najavits & Lester, 2008). This may be caused by the different cocaine effects on the brain in different sexes, which should be investigated in the future. Besides, the subject number of the original dataset is not large, which might introduce some bias to the results (Tommasi et al., 2017; Torralba & Efros, 2011). The representativeness of the results could be improved, and bias could be reduced if larger datasets from more diverse regions are available in the future. Also, the original dataset has more CUD subjects than the HC subjects. While the SVM is very sensitive to the imbalance of classes (Spelmen & Porkodi, 2018), this might cause a large difference between the sensitivity and specificity of results, which makes it important to keep the two classes balanced.

In conclusion, the present study supported the hypothesis that during the binge/intoxication stage of cocaine-use disorder, the Addictions Neuroclinical Assessment circuit is more consistent for analysis than the triple network model, as the default mode network is not as important as the other two networks. The machine learning classifier based on the salience network and the central executive network, which distinguishes CUD subjects from HC subjects, had the best performance. The significant relationship observed between the classification score of the ML model and measures of CUD patients, along with the functional connectivity changes between CUD patients and HC subjects, suggested the cumulative impact of cocaine use on brain function as the biomarker of cocaine severity.

## Acknowledgments

Research reported in this publication was supported by the National Cancer Institute of the National Institutes of Health under Award Number P30CA036727. This study was supported, in part, by the National Institute on Alcohol Abuse and Alcoholism (P50AA030407-5126, Pilot Core grant). This study was also supported by the Nebraska EPSCoR FIRST Award (OIA-2044049). This work was also partially supported by the National Institute of General Medical Sciences under Award Numbers P20GM103427, P20GM130447, and 1U54GM115458-01. This study was in part financially supported by the National Institute of Mental Health under Award Number 5U24MH100925. This work was also partially supported by the University of Nebraska Collaboration Initiative Grant from the Nebraska Research Initiative (NRI). This work was also partially supported by the Office of The Director, National Institutes of Health of the National Institutes of Health under Award Number R03OD038391. The content is solely the responsibility of the authors and does not necessarily represent the official views from the funding organizations.

## Notes

### Competing Interest Statement

The authors have declared no competing interest.

