## Supplementary material for "Functional Connectivity Alterations in Cocaine Use Disorder: Insights from the Triple Network Model and the Addictions Neuroclinical Assessment Framework": Table S1 and Table S2

Table S1 Altered Functional Connectivity within the DMN, SN, and CEN in CUD Subjects Compared to HC Subjects (Voxel p < 0.01, Cluster p < 0.05, Two-Tailed GRF Corrected; Anatomical Regions Identified by AAL Atlas)

| Type | Network | Anatomical Region | Cluster size | MNI coordinate (Peak) | | | T Values |
| --- | --- | --- | --- | --- | --- | --- | --- |
|  |  |  |  | **X** | **Y** | **Z** |  |
| CUD < HC | DMN | Right middle temporal gyrus | 391 | 48 | -21 | -9 | -5.05 |
|  |  | Left middle temporal gyrus | 128 | -48 | -36 | -6 | -4.07 |
|  |  | Right superior medial frontal gyrus | 109 | 9 | 57 | 21 | -4.43 |
|  |  | Left superior temporal pole | 104 | -54 | 9 | -15 | -4.51 |
|  | SN | Right supramarginal gyrus | 220 | 66 | -36 | 39 | -4.11 |
|  |  | Right superior temporal pole | 195 | 54 | 3 | -3 | -4.42 |
|  |  | Left insular cortex | 79 | -30 | 9 | 15 | -4.49 |
| CUD > HC | CEN | Right superior frontal gyrus | 79 | 21 | 48 | 30 | 4.16 |

CUD: Cocaine Use Disorder Patients, HC: Health Control Subjects

Table S2 Correlation coefficient between the classifier scores of the DMN/SN/CEN and clinical measures in CUD patients.

|  |  | Years of CUD | Days to the last usage | CUD onset age | Dose per week |
| --- | --- | --- | --- | --- | --- |
| DMN | r | 0.3517 | -0.4218 | -0.3779 | 0.1731 |
|  | p | 0.0575 | **0.0133** | **0.0380** | 0.3145 |
| SN | r | 0.1842 | -0.3605 | -0.1339 | 0.1424 |
|  | p | 0.3184 | **0.0350** | 0.4760 | 0.4133 |
| CEN | r | 0.2997 | 0.0398 | -0.3296 | 0.1588 |
|  | p | 0.0727 | 0.7996 | **0.0478** | 0.3610 |
| DMN-CEN | r | 0.4576 | -0.1106 | -0.4928 | 0.2701 |
|  | p | **0.0060** | 0.5271 | **0.0027** | 0.1093 |
| SN-DMN | r | 0.4202 | -0.3820 | -0.4034 | 0.1484 |
|  | p | **0.0128** | **0.0223** | **0.0172** | 0.4099 |
| SN-CEN | r | 0.3161 | -0.2410 | -0.2981 | 0.2974 |
|  | p | 0.0677 | 0.1591 | 0.0882 | 0.0742 |
| Triple network | r | 0.4720 | -0.3036 | -0.4783 | 0.2654 |
|  | p | **0.0039** | 0.0751 | **0.0036** | 0.1151 |
